# Glyco-engineered pentameric SARS-CoV-2 IgMs show superior activities compared to IgG1 orthologues

**DOI:** 10.1101/2022.12.22.521646

**Authors:** Somanath Kallolimath, Roman Palt, Esther Föderl-Höbenreich, Lin Sun, Qiang Chen, Florian Pruckner, Lukas Eidenberger, Richard Strasser, Kurt Zatloukal, Herta Steinkellner

## Abstract

Immunoglobulin M (IgM) is the largest antibody isotype with unique features like extensive glycosylation and oligomerization. Major hurdles in characterizing its properties are difficulties in the production of well-defined multimers. Here we report the expression of two SARS-CoV-2 neutralizing monoclonal antibodies in glycoengineered plants. Isotype switch from IgG1 to IgM resulted in the production of pentameric IgMs, comprising of correctly assembled 21 human protein subunits. All four recombinant monoclonal antibodies carried a highly reproducible human-type N-glycosylation profile, with a single dominant N-glycan species at each glycosite. Both pentameric IgMs exhibited increased antigen binding and virus neutralization potency, up to 390-fold, compared to the parental IgG1. Collectively, the results may impact on the future design of vaccines, diagnostics and antibody-based therapies and emphasize the versatile use of plants for the expression of highly complex human proteins with targeted posttranslational modifications.

## Introduction

Immunoglobulin M (IgM) is the largest antibody (Ab) isotype that is produced by the immune system of vertebrates. It can bind to a wide variety of pathogens and antigens (Ag) and is an essential part of the immune response. Comprehensive serological profiling in the course of diverse viral infections, including SARS-CoV-2, revealed a discerning appearance of various Ab iso- and subtypes, with increased levels of IgM, IgA, IgG1, and IgG3 (Amanat *et al*., 2020; Chakraborty *et al*., 2021; Gallerano *et al*., 2015). In fact, despite accounting for only ~12% of total immunoglobulins in plasma from healthy donors, IgG3 and IgM account for approximately 80% of the total neutralization in SARS-CoV-2 convalescent plasma (Kober *et al*., 2022). The overlapping or time-delayed response of antibody iso- and subtypes in infected individuals points to the versatile roles of Abs to combat infections effectively. Notwithstanding, research and industry have mainly focused on IgG1, with the consequence that the specific properties of other Ab isotypes are still poorly understood. This also applies to IgM Abs that bear interesting intrinsic features, like extensive N-glycosylation (more than 10% of the IgM molecular mass accounts for glycans) and circulate as ~950 kDa pentamers in human serum. In fact, the pentameric existence offers an avidity advantage and is a strong stimulus for the IgM typical complement activation (Polycarpou *et al*., 2020; Sharp *et al*., 2019).

Understanding the isotype-dependent properties requires the characterization of monoclonal antibodies (mAbs) of various isotypes against a defined antigen/epitope. However, the isolation of pathogen-specific monoclonal IgM is challenging due to low serum abundance, rapid decline as disease progress and class switch to various isotypes (Liu *et al*., 2020). Early studies of recombinantly produced human mAbs (Tiller *et al*., 2008) often involved isotype-switching of IgM to IgG1, mainly due to technical difficulties in the production of well-defined IgM pentamers. This approach has also been employed to study the properties of IgM Abs against SARS-CoV-2, however with inconclusive outcomes. (Wang *et al*., 2021) expressed a series of naturally selected anti-SARS-CoV-2 IgMs, switched to IgG1, thereby lowering the neutralizing (NT) activity. In another study, the complete loss of NT potency of two IgM Abs following isotype switch to IgG1 was reported (Callegari *et al*., 2022). Interestingly, isotype switch of two anti-influenza mAbs from IgM to IgG1 changed their neutralization potency. The activity of the more potent IgM was reduced by approximately 100-fold, while that of the less potent IgM did not change significantly (Shen *et al*., 2019). Although the vice versa engineering (i.e. IgG1 to IgM) often enhances potencies, this does not apply to all anti-SARS-CoV-2 Abs (Ku *et al*., 2021; Pisil *et al*., 2021). Notably, even Abs that bind to identical epitopes do not react equally. For example, the therapeutic anti-cancer IgG1 mAbs Pertuzumab and Trastuzumab, that bind to identical HER2 epitopes, behaved differently when switched to IgM (Samsudin *et al*., 2020). While Pertuzumab-IgM inhibited proliferation of HER2 over-expressing cells more effectively than its IgG1 counterpart, the reverse was observed for Trastuzumab. Taken together, previous observations highlight frequent uncertain consequences that accompany the class-switching of Abs and underline the importance of further research. Given the technical challenges in producing well-defined pentameric IgM molecules, direct comparisons of the functional attributes of IgM and IgG have been difficult.

Here we report the plant-based recombinant expression of two anti-SARS-CoV-2 mAb isotypes that share the same antigen-binding fragments (Fab). The original IgG1 mAbs with substantially different Ag-binding and virus NT properties were isotype-switched to IgM and produced as multimers. Biochemical and functional features of the IgG1 monomers and respective IgM pentamers, consisting of ten heavy and light chains (HC, LC) plus one joining chain (JC), were investigated. We demonstrate the correct assembly of recombinant mAbs and reveal superior Ag-binding and NT activities for the pentameric IgM isotypes when compared with their IgG1 orthologues. In addition, we elucidate the detailed N-glycosylation status of IgG1 and IgM mAbs by MS-based glycosite-specific profiling.

## Results and Discussion

### Production of recombinant monoclonal IgG1 and IgM

The two broadly neutralizing anti-SARS-CoV-2 mAbs P5C3 and H4 served as template in this study (Fenwick *et al*., 2021; Wu *et al*., 2020). Both Abs derive from convalescent human sera and are member of the IgG1 subclass. They bind to epitopes at the receptor-binding domain (RBD) of the spike protein, however their antigen-binding activities vary significantly. While P5C3 exhibits binding affinities in the picomolar range, these values are orders of magnitude higher for H4, depending on the virus isolate (Fenwick *et al*., 2021; Wu *et al*., 2020). To produce H4 and P5C3 in two different isotype formats (IgG1 and IgM), genes coding for corresponding heavy and light chain (HC, LC; Fig EV 1) were co-expressed in the glyco-engineered *Nicotiana benthamiana* line ΔXTFT (Strasser *et al*., 2008) through agroinfiltration. To facilitate pentamer formation, a gene coding for the joining chain (JC) was co-expressed with IgM. Four days post infiltration (dpi) recombinant IgMs and IgG1s were purified with affinity chromatography followed by size-exclusion chromatography (SEC). SDS-PAGE of the four purified mAbs confirmed the presence of the LC and HC without obvious degradation products or impurities (Figure 1A). Size exclusion chromatography (SEC) enabled the separation of IgM multimers from monomers (accounting for approximately 5-10% of purified IgM) and the presence of IgM pentamers was confirmed by SEC-MALS (Figure 1B). Collectively, four assembled mAb variants were generated: monomeric P5C3- and H4-IgG1; pentameric P5C3- and H4-IgM-P. Also, during purification minor amounts of monomeric P5C3- and H4-IgM-M were retrieved.

**Figure 1:**
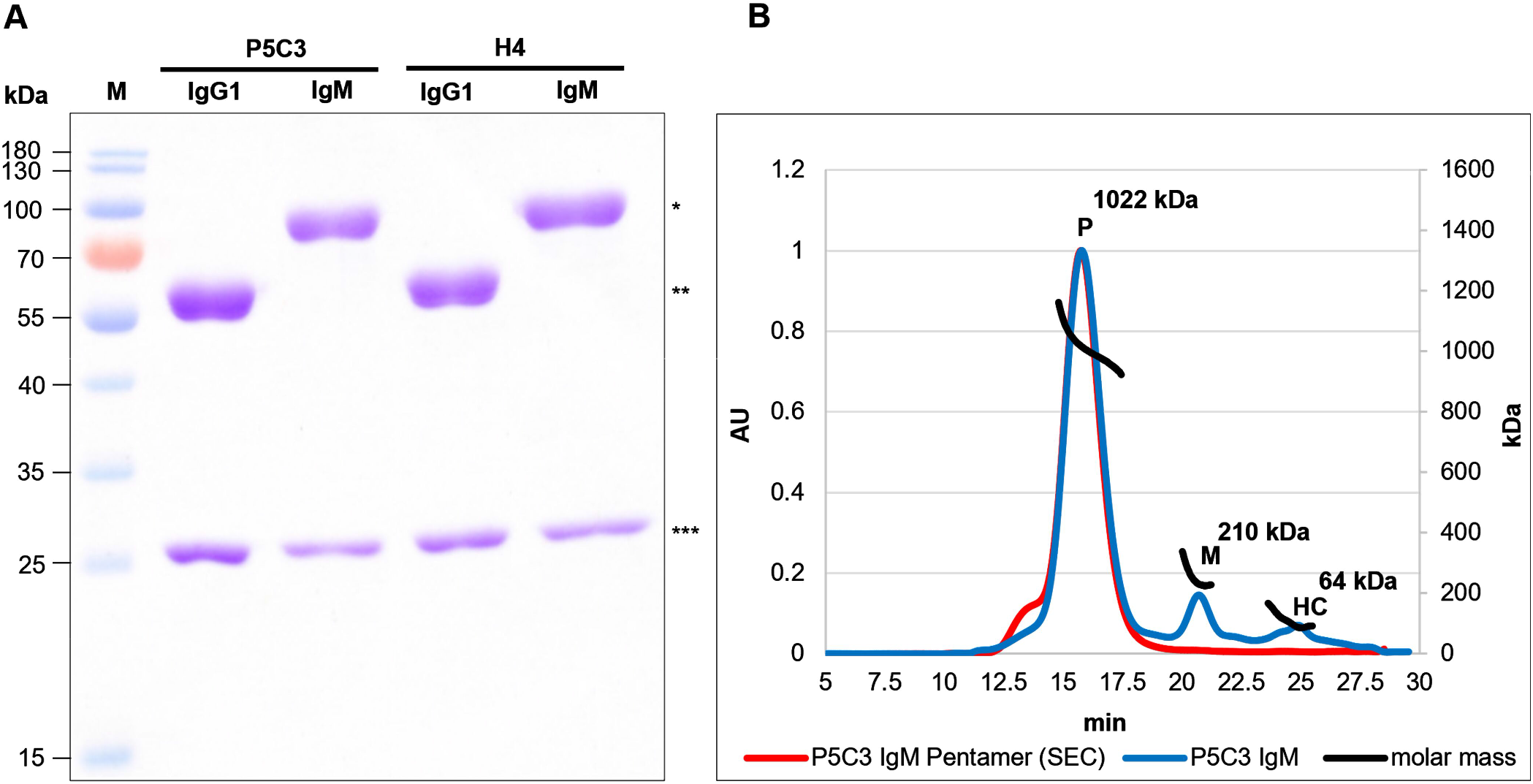
**A**: Reducing SDS-PAGE of P5C3- and H4-IgG1 and -IgM produced in ΔXTFT. 3 μg protein was loaded at each lane. M: Marker; * and **: IgM and IgG1 HC, respectively; ***: LC. **B:** SEC-MALS profiles of P5C3 IgM: Overlay of P5C3 IgM post affinity purification (blue line) and a SEC-isolated P5C3 IgM pentamer with minor amounts of higher multimers (red line); P5C3 IgM pentamers (~1022 kDa); monomers (~210 kDa); HC (~64 kDa); x-axis: retention time (in min), y-axis left: protein amount (arbitrary units), y axis-right: molecular mass (kDa).

### N-Glycosylation of plant-derived IgG1 and IgM

An interesting feature of IgM is its high N-glycosylation content, with five conserved N-glycosites (GS) at the HC and one at the JC. Overall, N-glycans account for more than 10% of the mAb’s molecular mass, with significant functional impacts (Colucci *et al*., 2015; Vattepu *et al*., 2022). GS 1-3 of pentameric human serum IgM (located in CH1-, CH2- and CH3-domains, respectively, Fig EV 2) carry complex sialylated structures. GS4 and 5 (located on the CH3 domain and the 18 amino acid-long C-terminal tailpiece, respectively) are decorated with oligomannosidic structures (Arnold *et al*., 2005; Loos *et al*., 2014). The single GS of the JC is highly sialylated. Whereas, IgG1 carries one conserved, Fc-located GS, usually decorated with N-acetylglucosamine (GlcNAc) or galactose-terminating complex N-glycans (Stadlmann *et al*., 2008). Glyco-engineered ΔXTFT line was used since it generates Abs with largely homogeneous and reproducible N-glycosylation profiles lacking plant specific residues (Jugler *et al*., 2022; Strasser *et al*., 2008). Moreover ΔXTFT-derived IgGs often exhibit increased functional activities compared to orthologues produced in CHO cells or wild type plants (Forthal *et al*., 2010; Loos *et al*., 2015; Zeitlin *et al*., 2011).

In order to determine the N-glycosylation status of ΔXTFT derived H4 and P5C3 mAbs, liquid chromatography-electrospray ionization-tandem mass spectrometry (LC-ESI-MS/MS) was performed. MS spectra of H4- and P5C3-IgG1 displayed a single dominant glycoform at the Fc GS, namely xylose and core fucose-free GlcNAc-terminated structures (predominantly GnGn), accompanied by ~8% mannosidic structures, as typical for ΔXTFT-produced IgGs (Strasser *et al*., 2008) (Figure 2, Table EV 1). MS spectra of IgMs were more diverse as exemplified in detail by P5C3-IgM. Both molecular forms (monomers and pentamers) were analysed separately. As expected, GS1-3 of P5C3-IgM-P are mainly decorated with complex N-glycan structures (89, 73 and 96 %, respectively) with some carrying core fucose (up to 15%). In addition, mannosidic glycans were detected (7, 23, 0%, respectively). GS1-3 of the monomeric pendant (P5C3-IgM-M) also carried complex N-glycans, however to a much lesser extend (64, 26 and 77 %) (Figure 2, Table EV 1). Interestingly, mannosidic structures increased (up to 70%) to the expense of complex N-glycans. IgM GS4 and 5 virtually exclusively carry oligomannosidic glycans (Man5 - Man9) in both molecular forms, in accordance with serum-isolated IgM (Arnold *et al*., 2005; Loos *et al*., 2015). The single conserved GS of the JC was decorated with 77% complex glycans, however, a significant proportion was incompletely processed (hybrid: Man4Gn, Man5Gn). Mannosidic structures accounted for approximately 23% (Figure 2, Table EV 1). While GS1, 2, and 4 were efficiently occupied, GS3 and 5 were glycosylated approx. 50% only (Table EV 1).

**Figure 2:**
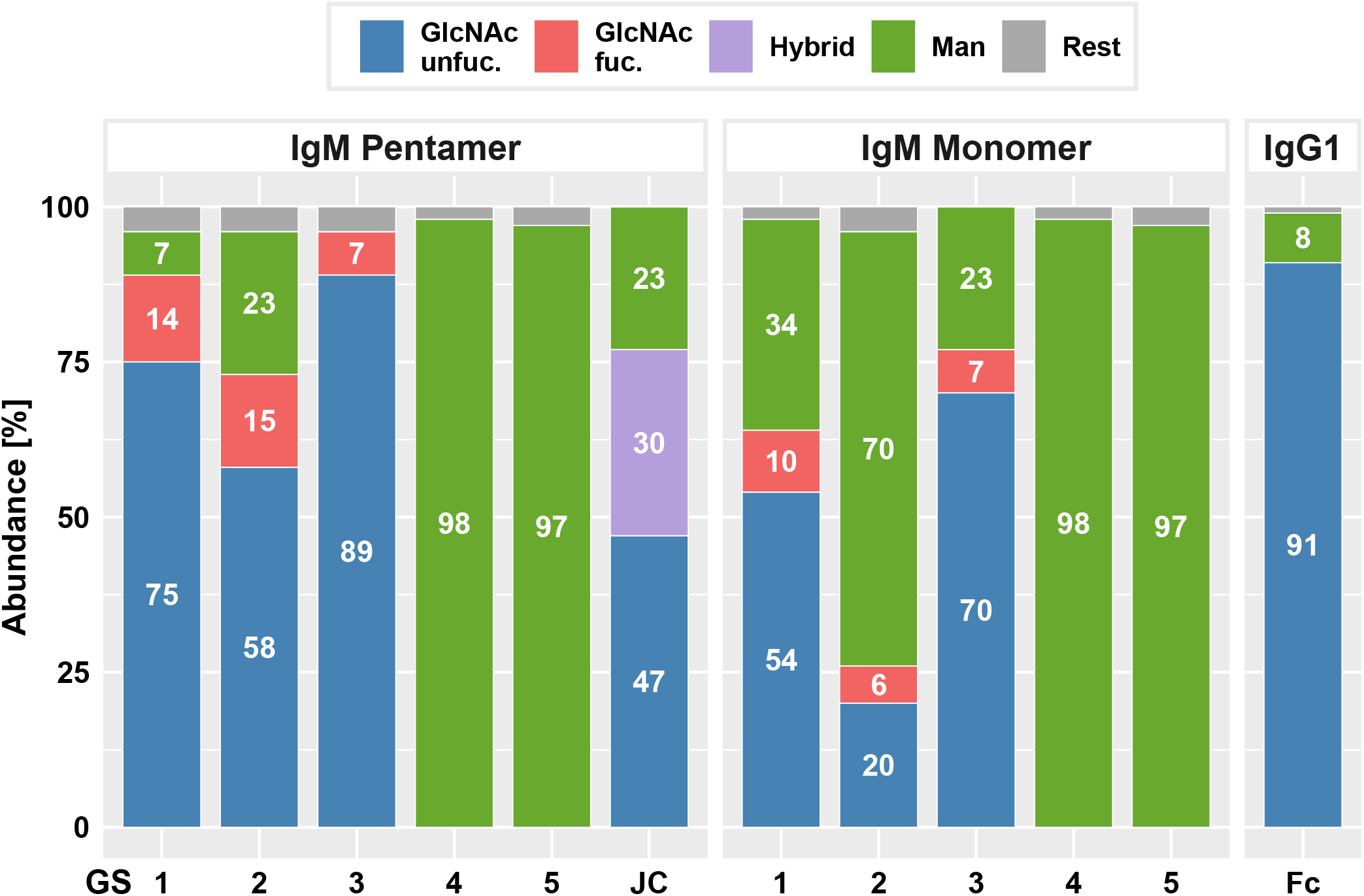
LC-ESI-MS/MS-derived N-glycosylation profiles of purified P5C3-IgG1, IgM penta- and monomers and joining chain (JC). Bars represent the relative abundance (%) of glycoforms present at each GS (for further details see EV table 1). Blue and red: non fucosylated and fucosylated complex GlcNAc-terminating N-glycans, respectively; purple: incompletely processed (hybrid) N-glycans; green: mannosidic N-glycans (Man5-Man9). Rest: combines detected glycans below 5%.

### Antigen binding assay using direct sandwich ELISA

Functional activity of anti-SARS-CoV-2 mAb variants were determined by antigen-binding assays. Direct sandwich ELISAs using SARS-CoV-2 spike protein RBD (Wuhan strain) as antigen and HRPO-labeled mAb CR3022 as secondary antibody was performed. Coating plates with target Abs directs the antigen to bind in a specific orientation, in contrast to direct coating of antigens that would result in random orientation, thereby reducing consistency. CR3022, initially developed against SARS-CoV, broadly detects SARS-related corona viruses (Yuan *et al*., 2022) and does not compete with binding of H4 and P5C3, respectively. First, the binding properties of the IgG1 variants (H4- and P5C3-IgG1) were determined (Figure 3A). As expected, P5C3-IgG1 exhibited a ~10-fold greater binding activity compared to H4-IgG1 (EC_50_ 106 pM for P5C3 and 1195 pM for H4). This feature also translated to the monomeric IgM-M (EC_50_ 90 and 974 pM for P5C3 and H4, respectively). In accordance, similar antigen-binding activities were observed when comparing corresponding IgG1 and IgM-M forms, i.e., P5C3-IgG1 versus IgM-M; and H4-IgG1 versus IgM-M (Figure 3A). By contrast to the monomeric IgG1 and IgM-M, IgM-Ps showed ~7-fold increased binding activities (EC_50_ of IgM-P of 16 and 169 pM for P5C3 and H4, respectively). Collectively, our results demonstrate the plant-based expression of glyco-engineered and isotype-switched functionally active P5C3 and H4 mAbs.

**Figure 3:**
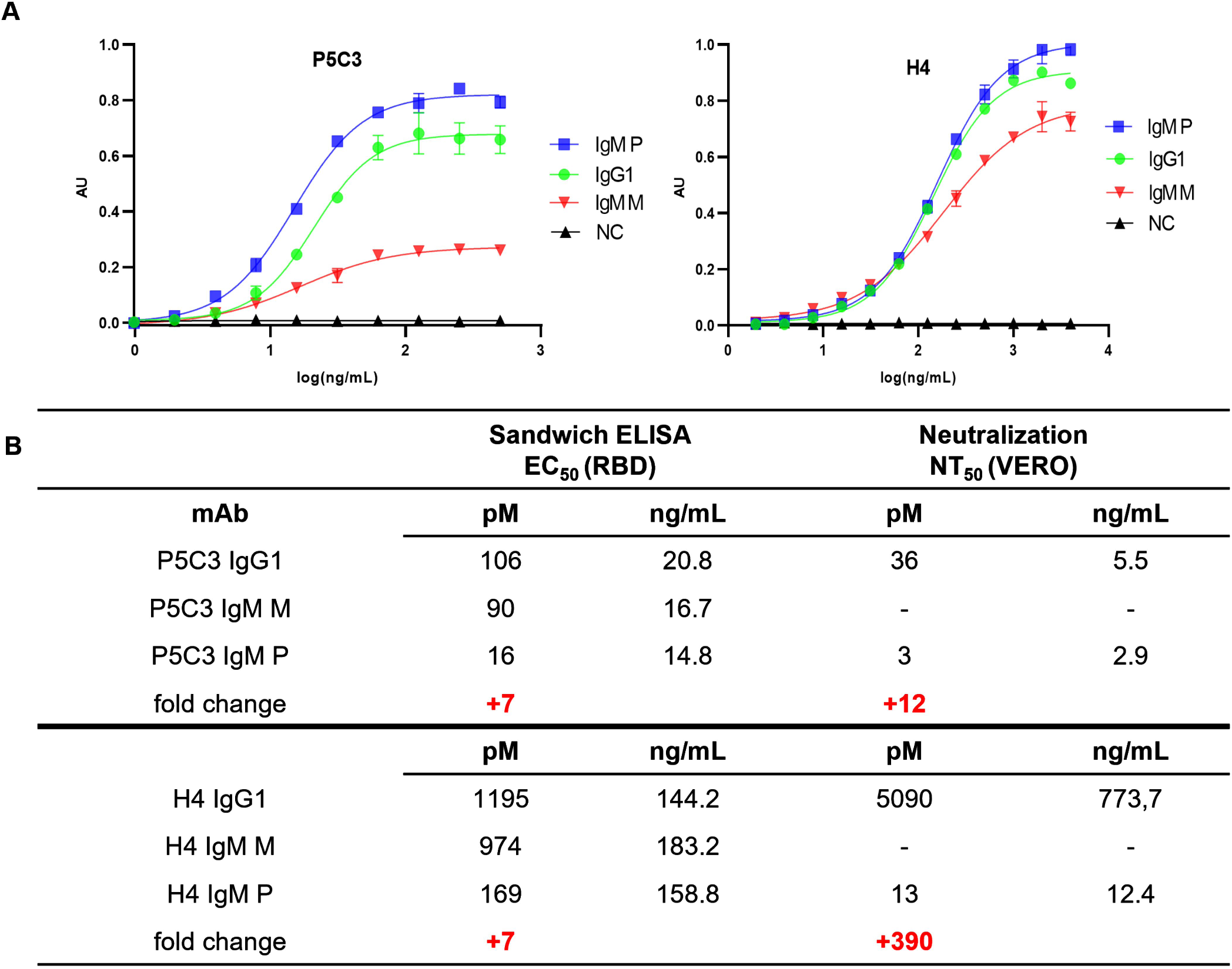
**A:** Sandwich ELISA: antigen-binding activity of purified mAbs (EC_50_ values in picomoles/L) to recombinant RBD, using CR3022-HRP for detection. X-axis: concentration (ng/mL); y-axis: absorbance (AU); **B:** Ag-binding EC_50_ and neutralization NT_50_ values. The fold change of EC_50_ and NT_50_ values between IgG1 and IgM-P is highlighted in red. IgM-M, IgM-P: IgM mono- and pentamers, respectively; NC: negative control (irrelevant IgG1 mAb).

### Neutralization activity of IgG1 and IgM pentamers

To determine neutralization activities of P5C3 and H4 Abs, Vero cell–based SARS-CoV-2 plaque reduction assays according to (Bewley *et al*., 2021) were performed. When comparing the two IgG1s, P5C3 showed an approximately 140 times higher virus NT potency than H4 (IC_50_ = 36 and 5090 pM for P5C3- and H4-IgG1, respectively; Figure 3B, Figure EV 3). This is in accordance with previous observations (Fenwick *et al*., 2021; Wu *et al*., 2020). Interestingly, when comparing the NT activities of IgG1s to the corresponding IgM-Ps, P5C3-IgM-P exhibited a ~12 times increase (IC_50_ 3 pM) and H4-IgM-P a ~390 times increase (IC_50_ 13 pM). Note, the remarkable high neutralization potency of P5C3-IgM-P that results in extremely low IC_50_ values might skew the results and it could well be that the activity difference of IgG1 to IgM-P is considerably higher for that mAb.

Collectively, the properties of IgM are only beginning to be explored in depth, which partly is due to the challenges in the generation of pentamers. Using a plant-based approach, we generated highly pure IgM pentamers. This was achieved by the simultaneous delivery of three expression constructs (HC, LC and JC) to plants and a subsequent two-step purification procedure.

While the production of IgG1 has been demonstrated in multiple cases using plant-based expression systems, only a few studies report the generation of more difficult to express Abs like IgG3 and dimeric IgA (Goritzer *et al*., 2020; Kallolimath *et al*., 2020; Kallolimath *et al*., 2021; Sun *et al*., 2021; Teh *et al*., 2021; Vasilev *et al*., 2016). Although an IgG1 to IgM switched mAb was produced recently (Jugler *et al*., 2022), only one previous report describes the plant-based production of multimeric IgM (Loos *et al*., 2014). The study presents a proof-of-concept for the multimeric assembly of an anti-cancer IgM in plants. The plant-derived IgM formed hexamers and pentamers in a 1:1 ratio, while the same IgM produced in a human cell line (PER.C6) assembled predominantly to pentamers. Here we obtained IgM pentamer formation of ~90% and anticipate protein-intrinsic factors that impact on specific multimer formation.

Importantly, using the glycoengineered ΔXTFT line (Strasser *et al*., 2008) enabled the generation of IgG1 and IgM mAbs with a largely homogenous N-glycosylation profile, largely devoid of plant specific glycosylation. Nevertheless, some glycans at GS 1-3 of IgM are fucosylated, a feature also observed for other glycoproteins expressed in this RNAi line (Gattinger *et al*., 2021). While each GS carried a highly reproducible single dominant N-glycan species, the pool of N-glycans in mammalian cell-produced mAbs is by far more heterogeneous, with substantial differences depending on the production conditions (Hennicke *et al*., 2017; Hennicke *et al*., 2019). Significantly, large glycan homogeneity enables the generation of well-defined mAbs, an important parameter not only to study the Ab properties but also to meet high biopharmaceutical (production) standards. Moreover, consistent N-glycosylation allows the engineering towards targeted structures in plants, like protein sialylation (Loos *et al*., 2014), facilitating the investigation of the so far largely unknown impact of this important post translational modification on the biological activities (Colucci *et al*., 2015). Interestingly, we found some differences in the N-glycosylation composition between penta- and monomeric IgM at GS1-3. This has not been reported so far, most probably due to the low abundance of this molecular form in mammalian cells. Nevertheless, a study that compared the N-glycosylation of penta- and hexameric IgMs produced in human cells revealed differences at GS1-3, mainly in branching and sialylation of complex N-glycans (Moh *et al*., 2016). However, whether such alterations have a structural and/or functional impact is not known. It is also remarkable that site specific N-glycosylation (complex and mannosidic N-glycans, respectively) of plant produced IgM matches that of the human serum Abs. Although the molecular mechanisms for site specific glycosylation are not well understood, they seem to be conserved across kingdoms. Notably, a series of studies demonstrated that ΔXTFT-produced mAbs against different human viruses exhibit increased functional activities compared to mammalian cell- or wild type plant-produced counterparts, in vitro and in vivo, e.g. (Forthal *et al*., 2010; He *et al*., 2014; Hurtado *et al*., 2020; Zeitlin *et al*., 2011). Whether this translates to anti-SARS-CoV-2 mAbs is currently not known, however there is strong evidence that the Ab glycosylation signature plays a critical role in the pathogenesis of COVID-19 (Chakraborty *et al*., 2021; Hoepel *et al*., 2021; Larsen *et al*., 2021; Siekman *et al*., 2022).

In this study, using sandwich ELISA, we demonstrate a sevenfold increased Ag-binding of H4- and P5C3-IgM-P to recombinant RBD, compared to their IgG1 orthologues. This is relatively modest when compared to other reports that demonstrated a more than 100-fold increase in RBD-binding by IgG1/IgM switch (Ku *et al*., 2021). On the other hand, a predictable enhancement in binding was not possible (Ku et al., 2021), indicating an epitope/paratope specific impact. It should be noted that methodological differences between studies, e.g., ELISA settings, might affect specific results. Nevertheless, there is mounting evidence that an isotype switch of IgG1 to IgM-P results in increased Ag-binding in many cases. This might have direct consequences on the development of improved diagnostics. While the highly sensitive nucleic acid-based tests are considered the golden standard for the detection of SARS-CoV-2 infections, they require specialized instruments and cannot easily be implemented as point of care (POC) diagnostics. Current SARS-CoV-2 Ag-POC tests use specific IgG1 mAbs in their settings, however in the light of the present (and related) results, pentameric IgMs might be a more suitable detection reagent.

Both IgM-P molecules exhibited substantially enhanced NT potency compared to the respective IgG1 orthologue, reaching a 390 times increase for H4. Notably, the 12 times NT enhancement of P5C3-IgM-P is most probably an underestimation as the extremely high potency of this mAb reached very low IC_50_ levels. Nevertheless, an Ag or Ab-specific effect as described previously cannot be excluded (Ku *et al*., 2021; Pisil *et al*., 2021; Shen *et al*., 2019). Our results are in agreement with recent investigations on anti-SARS-CoV-2 IgG3 and dimeric IgA mAbs (Kallolimath *et al*., 2021; Pisil *et al*., 2021; Sun *et al*., 2021; Wang *et al*., 2021) and with HIV data (Bournazos *et al*., 2016) which demonstrated significant NT improvements compared to IgG1. NT enhancements in orders of magnitude cannot exclusively be explained by the higher avidity of the multimeric structure and the results corroborate the bonus effect of Ab multi-valency that defines avidity not as the simple sum of individual binding site affinities (Freyn *et al*., 2021; Renegar *et al*., 1998). Superior NT potency of the IgM-P compared to modest variations in Ag-binding capacity of monomeric counterparts suggests that cross-linking the spike protein on the viral surface might be a critical factor. We hypothesize that cross-linking lowers the concentration of Abs required for NT and low spike densities facilitate Ab evasion. Interestingly, the spike protein copy number per SARS-CoV-2 virion is comparable with HIV, but 5 to 10 times less than that of other enveloped viruses, such as the influenza virus (Ke *et al*., 2020; Yao *et al*., 2020). In this context it is striking that, to our knowledge, such effective multimeric Ab induced NT enhancements have not been reported for e.g., influenza viruses. Another factor that could in part explain the gap between mAb binding- and NT is ACE2, which might impact on the NT assay. Furthermore, the observed effects of IgG1/IgM-P class switch are ascribed to factors like steric issues or altered epitope/paratope accessibility (Samsudin *et al*., 2020; Thouvenel *et al*., 2021). Also, given that Ab-Ag interactions can be drastically affected by small changes in different Ab domains (e.g. in light chain, hinge, V-region pairing, and VH and CH gene families, (Ling *et al*., 2018; Lua *et al*., 2019; Su *et al*., 2017; Su *et al*., 2018; Torres *et al*., 2005), it is necessary to investigate pentameric IgMs in a holistic approach. However, this provides challenges largely due to the experimental limitations associated with studying such large multimeric proteins. As a consequence, the molecular basis for how IgM achieves its strong avidity remains elusive, which hinders future development of therapeutic antibodies and vaccine design. The rapid and scalable expression of well-defined pentameric IgMs as shown here, may pave the way to overcome the current limitations and may contribute to further exploring of the properties of this highly interesting, but so far underexplored Ab isotype. The work underscores the versatile use of plants for the rapid expression and engineering of highly complex human proteins.

## MATERIALS AND METHODS

### Generation of P5C3-, H4-IgG1 and -IgM expression vectors

Codon-optimized HC and kappa (κ) LC variable fragment (Fv) sequences of P5C3 and H4 *P5C3-HCFv* (369 bp), *H4-HCFv* (379 bp); *P5C3-LCFv* (324 bp), *H4-LCFv* (339 bp) were grafted onto magnICON® vectors containing constant domains of human IgG1 HC, IgM HC and κLC by using BsaI restriction sites (Marillonnet *et al*., 2005), resulting in *P5C3-IgG1-HC* (1362 bp), *H4-IgG1-HC* (1371 bp) *P5C3-IgM-HC* (1731 bp), *H4-IgM-HC* (1741 bp) and the light chains *P5C3-κLC* (648 bp) and *H4-κLC* (663 bp). All vectors carry a barley α-amylase signal sequence for peptide secretion. Distribution among two compatible magnICON® plasmids was as follows: pICH26211: P5C3-IgG1, H4-IgG1, P5C3-IgM, H4-IgM; pICH31160: P5C3-IgGLC and H4-IgGLC (for vector details see Castilho & Steinkellner (2016)). Sequence information is available at supporting information. All constructs were transformed into Agrobacteria (strain GV3101 pMP90). The resulting strains were used for subsequent agroinfiltration experiments.

### In planta expression and purification of P5C3-, H4-IgG1 and -IgM mAbs

*Nicotiana benthamiana* plants (ΔXTFT line (Strasser *et al*., 2008)) were grown in a growth chamber under controlled conditions at 24 °C, 60% humidity with a 16 h light/8 h dark photoperiod. For agro-infiltration, respective recombinant bacterial strains were grown at 29 °C for 24 h, centrifuged at 4000 g for 10 min and resuspended in infiltration buffer (10 mM MES, pH 5.6; 10 mM MgSO_4_). Optical density of each strain was measured by extinction at 600 nm (OD_600_) of an adequate dilution. Final OD_600_ was set to 0.1 by dilution with infiltration buffer. Agroinfiltration mixes were delivered to leaves of 4-5 weeks old plants using a syringe. To produce H4- and P5C3-IgG1 isotypes, corresponding constructs carrying the heavy chains (P5C3-IgG1-HC or H4-IgG1-HC) and kappa light chains (P5C3-κLC or H4-κLC) were co-expressed. For the production of IgM agrobacteria carrying either P5C3-IgM-HC or H4-IgM-HC and corresponding light chain constructs (P5C3-κLC or H4-κLC) were coinfiltrated. Also, an agro-strain carrying the JC (Loos *et al*., 2014) was co-delivered. Infiltrated leaves were harvested 4 dpi, flash-frozen in liquid nitrogen, and ground to fine powder. Total soluble proteins (TSPs) were extracted with extraction buffer (0.5 M NaCl, 0.1 M Tris, 1 mM EDTA, 40 mM ascorbic acid; pH 7.4) in a ratio of 1:2 w/v (fresh leaf weight/buffer) for 90 min at 4 °C on an orbital shaker. Subsequently, the solution was centrifuged twice at 14,000 g for 20 min at 4°C and the supernatant vacuum filtrated using 8-12 μm and 2-3 μm filters (ROTILABO® Typ 12A and 15A).

Recombinant IgG1 was purified by affinity chromatography using protein A (rProA Amicogen, Cat no: 1080025), IgM purification was performed by POROS^™^ CaptureSelect^™^ IgM Affinity Matrix (Thermo Scientific^™^, Cat no: 1080025). TSP extracts were loaded at a flow rate of 1.5 mL/min on a manually packed column which was pre-equilibrated with 10 column volumes (CV) PBS (137 mM NaCl, 3 mM KCl, 10 mM Na_2_HPO_4_, 1.8 mM KH_2_PO_4_; pH 7.4). Washing was done with 20 CV PBS. Antibodies were eluted in 1 mL fractions with 0.1 M Glycine/HCl (pH 3.0), eluates were immediately neutralized with 1 M Tris (pH 9.0) and dialyzed overnight against PBS.

SEC-MALS was performed on a Shimadzu LC-20A Prominence system equipped with a diode array detector SPD M20A and a refractive index (RI) detector RID 20A. MALS data was acquired using a miniDAWN treos detector (Wyatt Technology, Santa Barbara, CA, USA). LabSolution Software (Shimadzu) and ASTRA V software were used for data collection. The samples were analyzed by using a dn/dc value of 0.185 mL/g as input for the MW calculation. MALS detector calibration was performed using BSA monomer (Merck) Monomeric and pentameric IgMs were separated by SEC. A Superose 6 Increase 10/300 GL, 10 mm i.d. × 300 mm column length (Cytiva Europe GmbH, Freiburg, Germany) column was used for the SEC experiments. Mobile-phase flow rate was set at 0.75 mL/min. A 60 min isocratic analysis was performed using PBS containing 0.2 M sodium chloride as mobile phase. For analysis 210-240 μg protein at about 1 mg/ml was loaded. Column calibration was performed with a set of molecular mass standards ranging from 1.3 to 670 kDa (Bio-RAD).

The fractions corresponding to the monomeric IgG1 and mono- and pentameric IgM were collected and concentrated with Amicon centrifugal filters, MWCO 10,000 kDa (Merck Millipore, UFC5010). SDS-PAGE analyses were performed in 12% gels under conditions. Gels were stained with Coomassie Brilliant Blue R 250 staining (Carl Roth GmbH + Co. KG). Concentrations were determined by spectrophotometer (NanoDrop^™^ 2000, Thermo Scientific). All purifications were performed at 4 °C.

### N-Glycan Analysis

The N-glycosylation profiles of the purified Abs were determined by mass spectrometry (MS) as described previously (Kallolimath *et al*., 2021; Sun *et al*., 2021). Briefly, respective heavy chains were excised from an SDS-PA-gel, digested with trypsin for IgG and trypsin and Glu C for IgM, and analyzed with an LC-ESI-MS system (Thermo Orbitrap Exploris 480). The possible glycopeptides were identified as sets of peaks consisting of the peptide moiety and the attached N-glycan varying in the number of HexNAc units, hexose, deoxyhexose, and pentose residues. Manual glycopeptide searches were performed using FreeStyle 1.8 (Thermo), deconvolution was done using the extract function. The peak heights roughly reflect the molar ratios of the glycoforms. Nomenclature according to Consortium for Functional Glycomics (http://www.functionalglycomics.org) was used.

For peptide mapping the files were analysed using PEAKS (Bioinformatics Solutions Inc, Canada), which is suitable for performing MS/MS ion searches.

### Direct Sandwich ELISA

Purified mAbs were diluted with PBS to 0.5 μg/mL (P5C3) and 2.0 μg/mL (H4). Certain mAbs were loaded with 50 μL/well to 96 well microplates (Thermo fisher maxisorp, catlog No: M9410-1CS) and incubated overnight After three washes with PBS-T (PBS with 0.05% Tween 20), the plates were blocked with 3% fat free milk powder, dissolved in PBS-T, for 1.5 h at RT. Recombinant RBD (Wuhan strain, (Shin *et al*., 2021)) was diluted in blocking solution and applied to the coated plates in two-fold serial dilutions starting from 500 ng/mL with P5C3 and from 4 μg/mL with H4, respectively. After 2 h incubation at RT, washing steps were performed. For detection of bound RBD, 50 μL of anti-RBD mAb CR3022 conjugated with horseradish peroxidase was diluted 1:15,000 in blocking solution and plates were incubated with it for 1 h at RT. Detection was performed with 50 μL per well 3,3’,5,5’- tetramethylbenzidine (Thermo Fisher, J61325.AU), the reaction was stopped with 2 M H_2_SO_4_ after 5-7 minutes incubation. Absorbance was measured at 450 nm (reference 620 nm) using a Tecan Spark® spectrophotometer. All samples were analyzed at least in two technical replicates. EC_50_ values were calculated by non-linear regression of the blank-corrected data points based on a four-parametric log model with GraphPad Prism (version 9).

### SARS-CoV-2 Neutralization Test

Neutralization assays were performed according to (Bewley *et al*., 2021). Briefly, VeroE6 cells (VC-FTV6, Biomedica, Vienna, Austria) were seeded in 48-well plates to achieve 100% confluency on the of infection. mAb were serially diluted (2-fold) and incubated with SARS-CoV-2 (Delta variant GK/478K.V1 (B.1.617. 2+AY.x), GISAID name: hCoV-19/Austria/Graz-MUG21/2021) for 30 min at 37 °C. Two wells were infected with the same mAb/SARS-CoV-2 mixture or SARS-CoV-2 without mAb treatment (positive control) and incubated for 1 h at 37 °C. Subsequently, the inoculum was removed, and cells were overlayed with 1.5% carboxymethyl cellulose (Sigma-Aldrich, St. Louis, MO, USA). After 48 h, cells were fixed with 4% neutral buffered formalin and stained immunohistochemically as described previously (Hardt *et al*., 2022). The number of plaques counted for the positive control were set to 100%. To calculate the half maximal inhibitory concentration (IC_50_), normalized data were used for nonlinear regression analysis with variable slopes (GraphPad PRISM Version 9). All experimental procedures involving SARS-CoV-2 were performed in a BSL-3 laboratory.

## Acknowledgements

magnICON® vectors were thankfully provided by Victor Klimyuk (Icon Genetics GmbH). Mass spectrometry measurements were performed by Clemens Grünwald-Gruber and Daniel Maresch (Core Facility Mass Spectrometry, University of Natural Resources and Life Sciences, Vienna, Austria). This work was supported by the projects of the Austrian Science Fund appointed to HS (grants I 4328-B and I 3721-B30) and Doctoral Program Biomolecular Technology of Proteins (W 1224).

## Author contributions

Somanath Kallolimath: conceptualization; data curation; formal analysis; investigation; supervision; Roman Palt: data curation; formal analysis; investigation; Esther Föderl-Höbenreich: data curation; formal analysis; investigation; Lin Sun: data curation; formal analysis; Qiang Chen, data curation, provided material; Florian Pruckner data curation; formal analysis; Lukas Eidenberger: data curation; formal analysis; Richard Strasser: data curation, provided material; Kurt Zatloukal: data curation; supervision; funding acquisition; Steinkellner Herta: conceptualization; investigation; supervision; funding acquisition All: writing–original draft; writing–review and editing

**Expanded View Figure 1:** Peptide sequences of P5C3- and H4-IgG1, -IgM heavy chains, joining chain (JC, P5C3- and H4-κLC kappa light chains. Variable domain is underlined, conserved glycosites are indicated in red.

**Expanded View Figure 2: A:** Schematic presentation of human IgG1, IgM and J-chain, including their serum glycosylation status. Dots represent glycosylation sites (GS); numbering of IgG1 and J-chain refers to GS position; **B:** N-glycan symbols according to Consortium for Functional Glycomics (http://www.functionalglycomics.org/).

**Expanded View Figure 3:** mAbs were evaluated for neutralization potency against SARS-CoV-2 infection measured by plaque reduction assay. Shown values correspond to a representative experiment (each concentration was tested in duplicates).

**Expanded View Table 1:** Detailed description of site-specific N-glycan composition of different mAbs and JC. Number represent abundance in %; nonglyco refers to % of non-occupied glyco-site. N-glycan symbols according to Consortium for Functional Glycomics (http://www.functionalglycomics.org/).

